# Coarse Raman and optical diffraction tomographic imaging enable label-free phenotyping of isogenic breast cancer cells of varying metastatic potential

**DOI:** 10.1101/2020.09.23.309138

**Authors:** Santosh Kumar Paidi, Vaani Shah, Piyush Raj, Kristine Glunde, Rishikesh Pandey, Ishan Barman

**Author notes:** Both authors contributed equally to this work (Santosh Kumar Paidi, Vaani Shah). Corresponding author: Ishan Barman.

## Abstract

Identification of the metastatic potential represents one of the most important tasks for molecular imaging of cancer. While molecular imaging of metastases has witnessed substantial progress as an area of clinical inquiry, determining precisely what differentiates the metastatic phenotype has proven to be more elusive underscoring the need to marry emerging imaging techniques with tumor biology. In this study, we utilize both the morphological and molecular information provided by 3D optical diffraction tomography and Raman spectroscopy, respectively, to propose a label-free route for optical phenotyping of cancer cells at single-cell resolution. By using an isogenic panel of cell lines derived from MDA-MB-231 breast cancer cells that vary in their metastatic potential, we show that 3D refractive index tomograms can capture subtle morphological differences among the parental, circulating tumor cells, and lung metastatic cells. By leveraging the molecular specificity of Raman spectroscopy, we demonstrate that coarse Raman microscopy is capable of rapidly mapping a sufficient number of cells for training a random forest classifier that can accurately predict the metastatic potential of cells at a single-cell level. We also leverage multivariate curve resolution – alternating least squares decomposition of the spectral dataset to demarcate spectra from cytoplasm and nucleus, and test the feasibility of identifying metastatic phenotypes using the spectra only from the cytoplasmic and nuclear regions. Overall, our study provides a rationale for employing coarse Raman mapping to substantially reduce measurement time thereby enabling the acquisition of reasonably large training datasets that hold the key for label-free single-cell analysis and, consequently, for differentiation of indolent from aggressive phenotypes.

## Introduction

Timely assessment of risk is critical for the detection and treatment of metastatic disease, which remains the main reason for cancer-related mortality [1]. The current clinical standard for assessment of metastatic risk relies on pathologic examination of sentinel lymph nodes following biopsy. In addition to being an invasive procedure, identification of metastatic cells in lymph node biopsies can be challenging and lead to an increase in false negatives [2]. Early detection of metastasis requires tools that can recognize the metastatic potential of cancer cells derived from the primary tumor or liquid biopsies. As primary tumors grow, a small fraction of the cancer cells termed circulating tumor cells (CTC) undergo epithelial to mesenchymal transition (EMT), locally invade the surrounding stroma, intravasate and are shed into the bloodstream leveraging their enhanced motility [3]. A few of these cells survive in the circulation to extravasate, locally invade and form premetastatic niches in secondary organs (e.g. lungs in breast cancer), and colonize through re-acquisition of epithelial characteristics via mesenchymal to epithelial transition (MET) [3]. While our understanding of the processes involved in metastasis has improved substantially in recent years, detecting phenotypic subtypes with metastatic competence has proven to be elusive due in part to the substantial heterogeneity observed in these cell populations [4]. Furthermore, our understanding of what imparts metastatic potential remains rudimentary, and biomarkers that can recognize such competence across different carcinomas are still lacking.

Early genomic analyses of tumors revealed additional organ-specific mutations in metastatic tumors despite sharing common ancestors [5]. Similarly, transcriptional analyses of breast cancer metastasis to various organs including lungs and brain have also identified largely distinct signatures characteristic of organotropism [6, 7]. However, these population-based analyses require elaborate sample preparation and fail to capture the variations in phenotypes at a single-cell level. Recently, we and others have also investigated the physical properties associated with the differences in metastatic phenotypes in specialized microfluidic platforms [8-15]. The pursuit of isolating CTC from blood to determine the course of metastatic disease has resulted in the proliferation of several cell labeling methods that leverage known epithelial markers for identification [16-21]. However, the use of epithelial markers may not be sufficient to detect CTC that undergo EMT to acquire mesenchymal properties, particularly in triple-negative breast cancers [22]. Also, the sensitivity of these methods is challenged by the small number of known markers of metastatic progression that can be targeted simultaneously.

To address these challenges, several techniques based on optical microscopy and imaging have attempted label-free phenotyping of cancer cells [23-27]. For example, phenotypic changes of 4T1 murine breast cancer cells in response to drug treatment were characterized in terms of morphological parameters extracted from fluorescence images in 3D cultures [25]. Rohde and co-workers have developed an automated platform for morphological analysis of cellular phenotypes using transport-based morphometry [24], which we recently used to analyze the quantitative phase images of activated and naïve CD8^+^ T cells [28]. Similar optical methods have also been leveraged for single-cell analysis of cancer phenotypes [29, 30], which often require large datasets for building robust prediction models.

In this study, we employed 3D optical diffraction tomography (ODT) and label-free Raman spectroscopy to quantitatively investigate both morphological and molecular differences between isogenic breast cancer cells of varying metastatic potential. We used a set of three isogenic cell lines composed of the parental MDA-MB-231 triple-negative breast cancer cell line (P231), circulating tumor cells (CTC), and lung metastatic cells (LM) where the latter two were derived respectively from the circulation and lungs of a mouse bearing parental P231 cells [31, 32]. Compared to the widely used qualitative phase imaging methods, such as Zernike phase contrast and differential interference contrast, quantitative phase imaging methods recover the phase delay caused by the sample, decoupled from absorption information. ODT is a form of quantitative phase imaging that allows morphological analysis of single-cells based on their 3D refractive index (RI) profiles [33-35]. In addition to providing traditional measures of morphology such as area and aspect ratio, the RI information allows a label-free and non-contact route for the determination of cell dry mass and local thickness of specimens with nanometric sensitivity [36]. The additional morphological insights provided by optical diffraction tomography have been increasingly exploited for label-free and stain-free *in vitro* analysis of cells and tissues [33, 34, 37]. Yet, most of the cellular studies have focused on either visualization of morphological dynamics in response to external stimuli such as drug exposure in single-cells or rapid identification of cells such as bacteria and white blood cells using deep learning by leveraging large datasets [38, 39]. Its utility in assessment of phenotypic differences among closely related mammalian cancer cells, particularly in data-limited settings, remains largely unexplored.

Raman spectroscopy, on the other hand, provides a label-free route for assessment of biological specimens with exquisite molecular specificity [40, 41]. This optical technique, based on the inelastic scattering of light, probes vibrational modes of molecules and allows direct profiling of molecular composition of biological specimens including live cells and tissues in their native states [40-42]. The simple integration of Raman spectroscopy with optical microscopy facilitates seamless vibrational spectroscopic imaging at diffraction-limited spatial resolution with subcellular resolution. Several groups including our own laboratory have exploited the high resolution and rich molecular information afforded by Raman spectroscopic imaging to study the molecular progression of cancer [23, 41, 43-45]. Due to the low likelihood of spontaneous Raman scattering, most single-cell imaging studies have focused on employing nanoparticles for plasmonic enhancement of signals and selective tagging of subcellular regions of interest [46, 47]. Therefore, only a few studies have attempted label-free characterization of cells for studying biological processes associated with physiological changes, disease progression, and drug response [48-50]. Our recent label-free Raman investigation in pellets of isogenic breast cancer cell lines that exhibit organotropism to brain, liver, lung, and spine revealed distinct metastatic organ-specific spectral signatures that were confirmed by metabolomics analysis [23]. Due to the long acquisition time, label-free Raman spectroscopic studies have either exploited high-resolution single-cell maps for analysis of limited cells or bulk sampling of cell populations that permits the use of machine learning algorithms for classification problems by generating large datasets at the cost of spatial information. This tradeoff between obtaining higher spatial resolution maps and acquiring sufficiently large datasets amenable for machine learning has largely prevented the use of machine learning techniques to learn and predict cellular phenotypes from Raman images with single-cell analytical resolution.

Therefore, in this study, we sought to test whether morphological attributes encoded by ODT and biomolecular insights obtained using Raman spectroscopy can predict the phenotype of the closely related isogenic cell lines P231, CTC and LM of varying metastatic potentials with statistical confidence. Using the 3D RI profiles obtained from ODT of single-cells, we compared the distributions of morphological parameters such as area, aspect ratio, and dry mass across the three classes, and used their combination to train and test random forest classifiers for automated identification. By leveraging coarse Raman sampling of single cancer cells to reduce the acquisition time and obtain spectral maps from a larger number of cells, we explored the intersection of the abovementioned resolution-sampling tradeoff to find a solution for identifying metastatic phenotype of cancer cells with single-cell analytical resolution. To show that coarse Raman maps capture sufficient information for achieving single-cell phenotyping, we used random forests to iteratively test spectral maps of individual cells against classifiers trained on the data from the remaining cells in the dataset. Furthermore, we used multivariate curve resolution alternating least squares (MCR-ALS) analysis to identify the spectra from subcellular compartments and test the utility of random forest classification in predicting the metastatic phenotype when only spectra from either nucleus or cytoplasm are available. The ability to use specific subcellular regions of the cell for chemical imaging is expected to further reduce the spectral acquisition time and boost the number of cells in the training dataset to capture population heterogeneity. Such label-free identification of cells with high metastatic competence would not only have a profound impact on the prediction of a patient’s risk of developing metastasis but also inform the design of optimal, personalized therapeutic treatments.

## Materials and methods

### Cell culture

An isogenic panel of varying metastatic potential derived from the human breast cancer cell line MDA-MB-231 was used in this study. In addition to the td-Tomato expressing parental MDA-MB-231 cells (P231), the panel consisted of CTC and LM cells previously obtained after orthotopic implantation of the parental cells in the fourth right mammary fat pad of female athymic nu/nu female mouse (NCI) as detailed in our previous publications [31, 32]. The three cell lines were cultured in RPMI-1640 media supplemented with 10% fetal bovine serum (FBS), 100 U/ml penicillin, and 100 µg/ml streptomycin and maintained at 37 °C and 5% CO_2_ in a humidified incubator.

### Optical diffraction tomography and data analysis

The three cell lines were seeded in glass coverslip-bottom Petri dishes for tomography. The morphological assessment of the cells was performed on an ODT system (HT‐1H, Tomocube Inc., Republic of Korea) comprised of a 60X water-immersion objective (1.2 NA), an off-axis Mach-Zehnder interferometer with a 532 nm laser and a digital micromirror device (DMD) for tomographic scanning of each cell [51]. The 3-D RI distribution of the cells was reconstructed from the interferograms using the Fourier diffraction theorem as described previously [52]. TomoStuido (Tomocube Inc, Republic of Korea) was used to perform reconstruction and visualization of 3D RI maps and their 2D maximum intensity projections (MIP). The 2D MIP images were segmented using CellProfiler™ (v3.1.9) software to isolate single-cells using Otsu two-class thresholding and neglecting the partial cells at the boundaries of raw images [53, 54]. After segmentation, we obtained 57, 35, and 44 cells in P231, CTC, and LM classes respectively. The area of each cell was calculated by counting the number of non-zero pixels in their corresponding segmentation masks generated by the CellProfiler™ software [55]. Similarly, the perimeter and aspect ratio were calculated respectively as the number of non-zero pixels at the edges of the masks and the ratio of major and minor axes lengths [55]. The cell dry mass was calculated from the 3D RI profile [56]. The morphological parameters were used to train a random forest classifier with 100 trees using the MATLAB TreeBagger class and inspect the out-of-bag-error.

### Raman spectroscopic imaging

The cells from three different passages (biological repeats) for each class were seeded on quartz slides (1 in x 1 in) coated with poly-lysine and incubated overnight to facilitate cell attachment for Raman imaging. The cells were fixed using 4% paraformaldehyde and washed prior to imaging in phosphate buffered saline (PBS) at room temperature. Five cells from each slide (technical repeats) were randomly selected for Raman mapping. The coarse single-cell Raman imaging experiments were performed on a HORIBA XploRA PLUS confocal Raman microscope. A 532 nm diode laser was used for excitation and delivered to the sample via a 60X water immersion objective (1.2 NA). The backscattered Raman light was dispersed using an 1800 lines/mm diffraction grating and imaged on a thermoelectrically cooled CCD coupled to the microscope. The spectra were acquired from the points on a coarse rectangular grid overlaid on each single cell to obtain spectra from various subcellular regions and capture intracellular spatial heterogeneity. Each spectrum in the fingerprint region (600-1950 cm^-1^) was acquired by exposing the sample to a laser power *ca.* 1 mW at each point for 2.5s (5 accumulations of 0.5s exposure).

### Raman data analysis

All the Raman spectral analysis was performed in MATLAB 2017b (Mathworks) environment. Spectroscopic imaging of each cell provided a hyperspectral dataset, where each pixel on the rectangular mapping grid corresponds to a Raman spectrum. The hyperspectral datasets from all the cells were unfolded (by preserving the spatial information and cell identity) and concatenated to form a combined spectral dataset for further analysis. The spectra in the fingerprint region were subjected to background subtraction using a fifth-order best-fit polynomial-based fluorescence removal method and cosmic ray removal using median filtering on the groups of spectra from each cell. Next, the points on the mapping grid exterior of the imaged cell were identified by Otsu thresholding on the 1452 cm^-1^ peak (CH_2_ bending mode of proteins) intensity and labeled separately for further analysis. The spectra were finally vector normalized to remove the variations in laser power across the experiments.

To identify spectra from specific subcellular regions, we performed MCR-ALS analysis for decomposing each spectrum into its constituents by iterative fitting under nonnegativity constraints on the obtained component spectra (loadings) and their contributions (scores) [57]. The components were identified as rich in cytoplasm, nucleus, and background (quartz and water) characteristics and confirmed by re-constructing the score maps for each cell. Each spatial location in the cell is assigned either cytoplasm, nucleus, mixed or background based on the Otsu thresholding of the cytoplasm-like and nucleus-like component scores, which were both negatively correlated [54]. The scores corresponding to cytoplasm-like and nucleus-like constituents were compared across the three cell lines through violin plots with outlier suppression for clarity. The significance of differences between medians was determined using Wilcoxon rank-sum test with the conventional threshold.

Random forest classifiers (bootstrap-aggregated or bagged decision trees) were trained using the TreeBagger class in MATLAB to enable the identification of the metastatic phenotypes. We used a leave-one-cell-out protocol by leaving one cell out each time as a test case and training random forest classifiers on spectra from the remaining cells. Several iterations of training were performed for each test case by selecting randomized subsets of training data to ensure equal membership for all the three classes and to avoid overtraining for the class with high data availability. The spectra of the excluded cell were subjected as a test dataset and the class label for the entire cell was determined according to the following class assignment criterion. Since the test dataset (left-out cell) remained the same for all training iterations, the median of predicted labels for each spectrum was used for decision making at the cell level. For each cell, the majority class was assigned as the predicted class if its membership was at least 30% higher than the random chance prediction and 30% higher than the membership of the second majority class. If these conditions were not met by the majority class, the test cell was labeled unclassified. To verify the ability of cytoplasm and nucleus spectra for the identification of metastatic phenotype, the random forest classifiers were run on the cytoplasm- and nucleus-rich spectra identified by the MCR-ALS analysis, in addition to running them on the entire spectral dataset.

## Results and discussion

The availability of isogenic breast cancer cells of varying metastatic potential derived from the same MDA-MB-231 human breast cancer cell line (**Fig. 1A**) enabled us to investigate the utility of label-free optical imaging for the identification of metastatic phenotypes at the single-cell level. We used parental P231 cells along with their circulating (CTC) and lung metastatic (LM) variants to assess the efficacy of ODT (**Fig. 1B**) and Raman spectroscopy (**Fig. 1C**) for capturing the phenotypic differences in terms of their morphological and molecular attributes. Our previous characterization of these cell lines confirmed their distinct metastatic abilities commensurate with the stage and organ from which they were isolated [31]. Our recent investigation of the biophysical properties of these cells revealed that the LM cells are most motile and least stiff, which bestow them with unique invasive capability [8].

**Figure 1.**
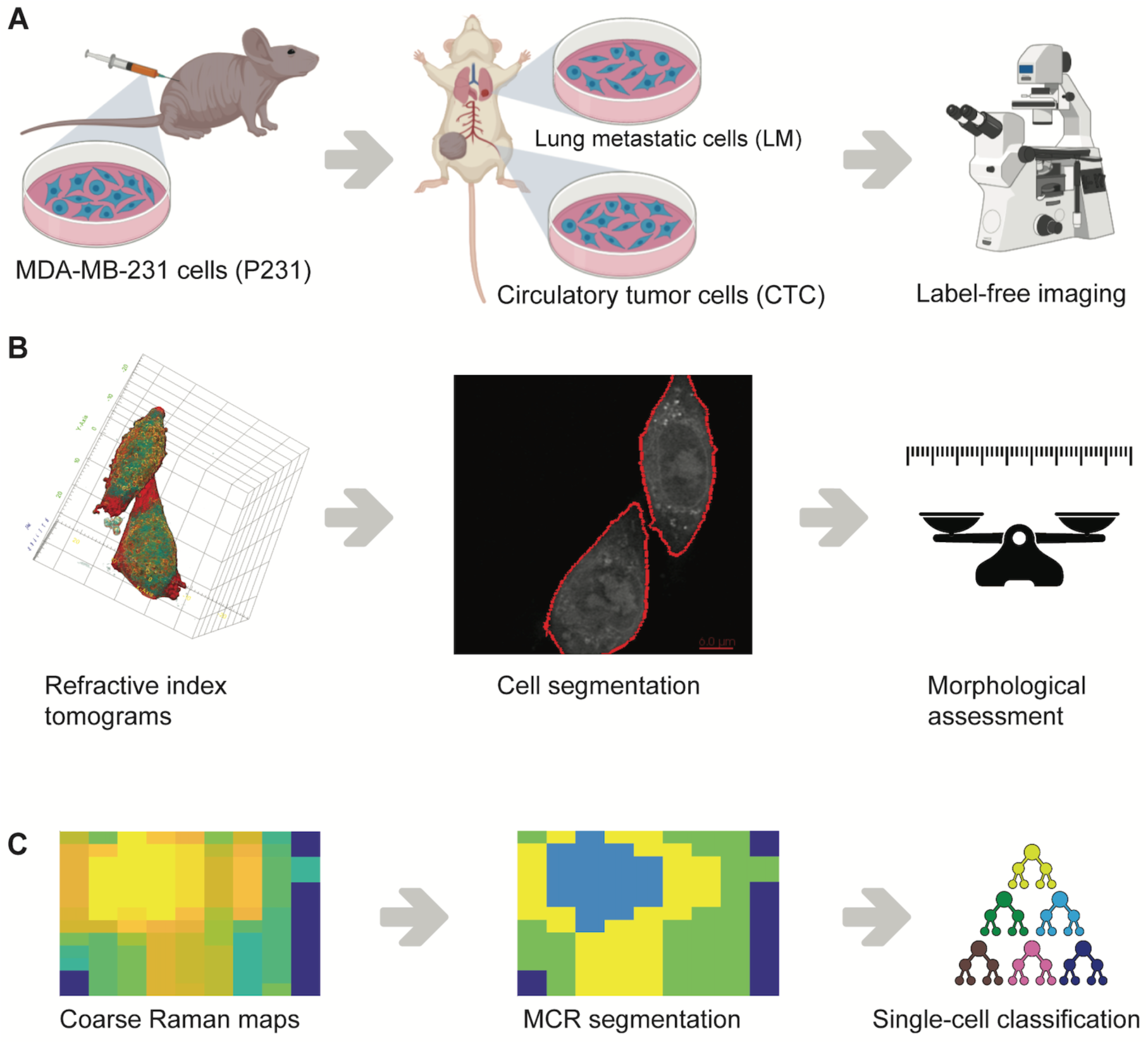
Label-free identification of metastatic phenotypes. **(A)** Circulating tumor cells (CTC) and lung metastatic cells (LM) used in the study were isolated from the blood and lungs of mice bearing parental MDA-MB-231 (P231) tumor xenografts. **(B)** Refractive index tomograms were segmented to isolate single-cells for morphological assessment. **(C)** Coarse Raman maps of single-cells were subjected to MCR-ALS analysis to identify subcellular regions rich in cytoplasm and nucleus prior to the use of supervised classification using random forests.

To understand the morphological differences among the phenotypically distinct isogenic cell lines, we acquired 3D RI tomograms of single cancer cells belonging to each group. While the RI tomograms of cells from all three cell groups (**Fig. 2A-C**) show expected intracellular heterogeneity arising from RI variations across subcellular compartments, the intercellular differences are not apparent from gross visual inspection. Therefore, the maximum intensity projections of the RI tomograms (**Fig. 2D-F**) were subjected to further assessment using CellProfiler™ software to quantify morphological parameters such as area and aspect ratio. Also, we calculated the cell dry mass directly from the 3D RI tomograms to include an additional dimension in the morphological analysis that cannot be readily measured from brightfield or phase contrast microscopy. As seen in **Fig. 2G**, we observed that the area of the cells increased steadily with the increase in metastatic potential from P231 to LM. However, a significant increase in aspect ratio (**Fig. 2H**) was only observed for the CTC in comparison to the P231 and LM classes, while the differences between the latter were not statistically significant. These observations are consistent with the characteristics of EMT and MET processes in metastasis of P231 cells to lungs that respectively result in the acquisition of a spindle shape by the CTC for enhanced motility to reach the metastatic site and re-acquisition of epithelial shape for promoting the proliferation of LM cells to form metastatic tumors [3]. We observed that the cell dry mass (**Fig. 2I**) increased with the metastatic potential of the isogenic cells, but the difference was statistically significant only for LM cells in comparison to P231 and CTC. The increase in the cell dry mass of the LM cells is consistent with the prior observation of an increased RI and cell dry mass of cancer cells in comparison with normal cells due to the higher accumulation of proteins associated with the higher proliferation of the former group [58]. Our observation expands this idea to the metastatic regime and provides a rationale to explore cell dry mass as a potential biomarker of invasiveness in future studies.

**Figure 2.**
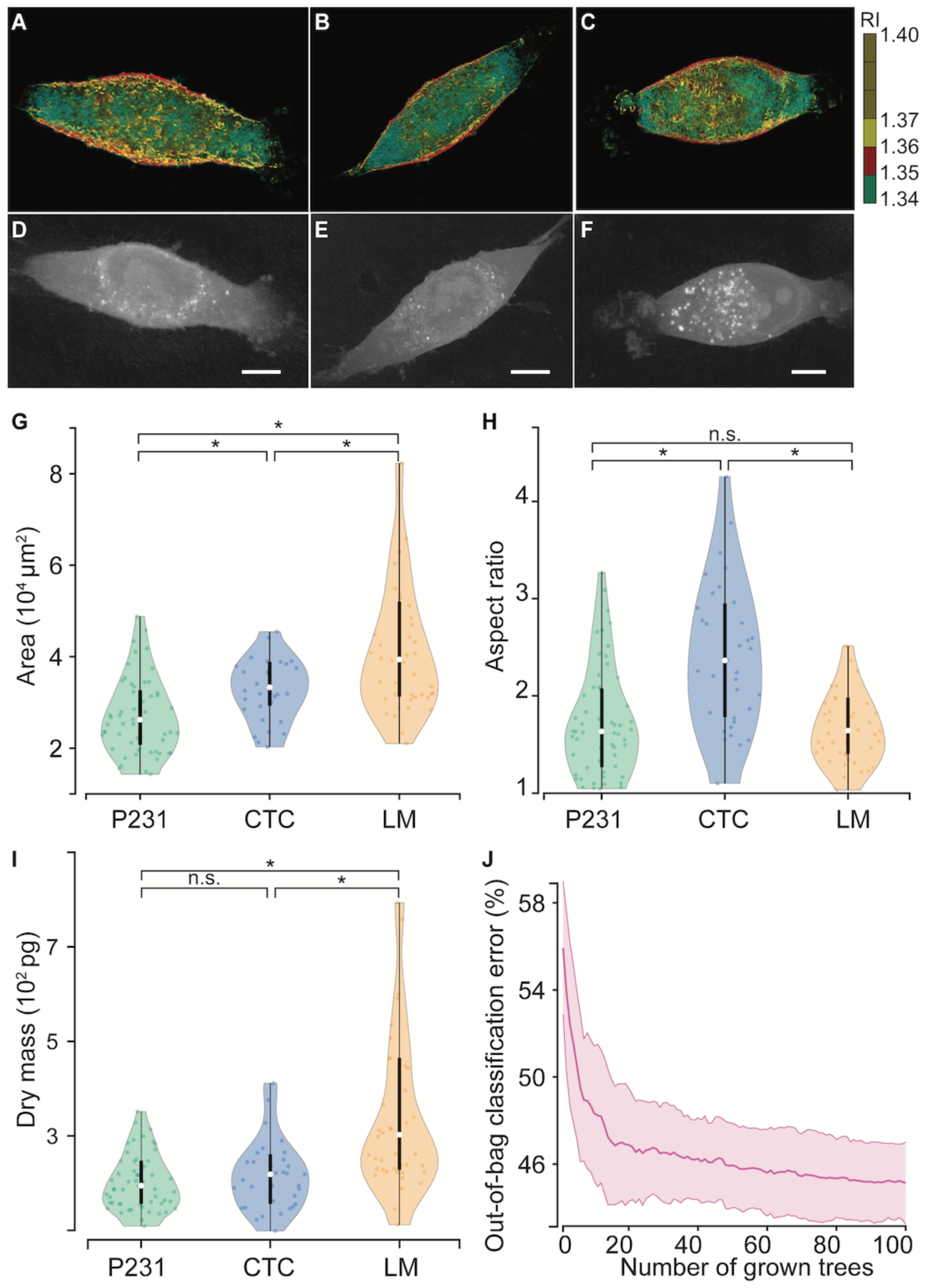
Morphological assessment of metastatic phenotypes. Representative 3D refractive tomograms of **(A)** P231, **(B)** CTC, and **(C)** LM cells show the intracellular variation of the refractive index. The maximum intensity projections of (**D**) P231, (**E**) CTC, and (**F**) LM were used for determining the 2D morphological parameters. The violin plots show the variations in **(G)** area, **(H)** aspect ratio, and **(I)** dry mass across the three classes. **(J)** The out-of-bag classification error plot shows that random forests build on the morphological parameters fail to accurately predict metastatic phenotypes. The scale bars represent 5 μm. * represents statistically significant differences at p < 0.05 threshold (Wilcoxon rank-sum test).

While the phenotypically distinct cell lines showed variable differences in individual morphological parameters, their utility for identifying phenotypes of single cancer cells is dependent on the existence of clear class boundaries between the three classes. The violin and box plots (**Fig. 2G-I**) show the appreciable overlap in the distribution of each morphological parameter across the metastatic potential thus making univariate analyses challenging for class separation. While prior studies have leveraged deep learning methods for cellular classification based on latent morphological features from the complete cell images, they have largely probed simpler systems such as bacteria, blood cells, immune cells, and anthrax spores compared to the current cohort of isogenic breast cancer cells [38, 39]. Since our current study is focused on the detection of the cellular phenotypes within the constraints of small training datasets, we trained random forest classifier to test if supervised models leveraging these three morphological parameters can accurately predict the metastatic phenotype of test samples. The out-of-bag classification error rate (**Fig. 2J**), calculated for each training sample by testing them against the decision trees in the forest that did not use them for training, was found to asymptotically plateau around 46% (compared to 33.3% random chance). These results indicate that while phenotype differences among the isogenic cells show subtle but significant morphological differences, they are not sufficient for robust classification of closely related cells at a single-cell analytical resolution, particularly when the training data is relatively scarce.

Next, we sought to check if molecular information provided by vibrational spectroscopy can enable the identification of metastatic phenotypes at a single-cell analytical resolution. Therefore, to build a dataset large enough for training machine learning models, we performed coarse Raman microscopy of all three isogenic cell lines. The entirety of each cell was mapped coarsely (average of 67 pixels per cell) to capture intracellular heterogeneity. While these coarse Raman images (**Fig. 3A**) do not offer diffraction-limited spatial resolution, they capture enough information for the cell classification task and help significantly reduce the spectral acquisition time for each cell. The mean (+/- 1 s.d.) of the spectra from the three cell lines (**Fig. 3B**) show prominent peaks at 931 cm^-1^, 1003 cm^-1^, 1085 cm^-1^, 1303 cm^-1^, 1450 cm^-1^, 1658 cm^-1^ indicative of the common biological constituents of cells and tissues [59]. Since there are no discernible visible differences between the spectra of the three cell lines, we used MCR-ALS analysis to decompose the spectra into component spectra and their scores. MCR-ALS decomposition allows representation of each spectrum in the dataset as a weighted sum of iteratively generated pure component-like basis spectra, without requiring any composition estimates as inputs [57]. In this study, a simple three-component MCR-ALS decomposition provided component loadings harboring features of cytoplasm, nucleus, and quartz background from the slide on which the cells were cultured (**Fig. 3C**). We identified MC1 and MC2 as loadings resembling cytoplasm and nucleus due to the prominence of cytoplasm features at 1003 cm^-1^ (C–C stretching vibration of the aromatic ring in the phenylalanine side chain) and 1639 cm^-1^ (amide I feature in proteins) in the former and nucleic acid features at 788 cm^-1^ (O-P-O stretching in DNA) and 1092 cm^-1^ (symmetric PO_2_^-^ stretching in DNA) in the latter. The assignment is also justified by the strong negative correlation between the MC1 and MC2 scores for each cell, assessed by an average correlation coefficient of −0.98 over all the cells in the study.

**Figure 3.**
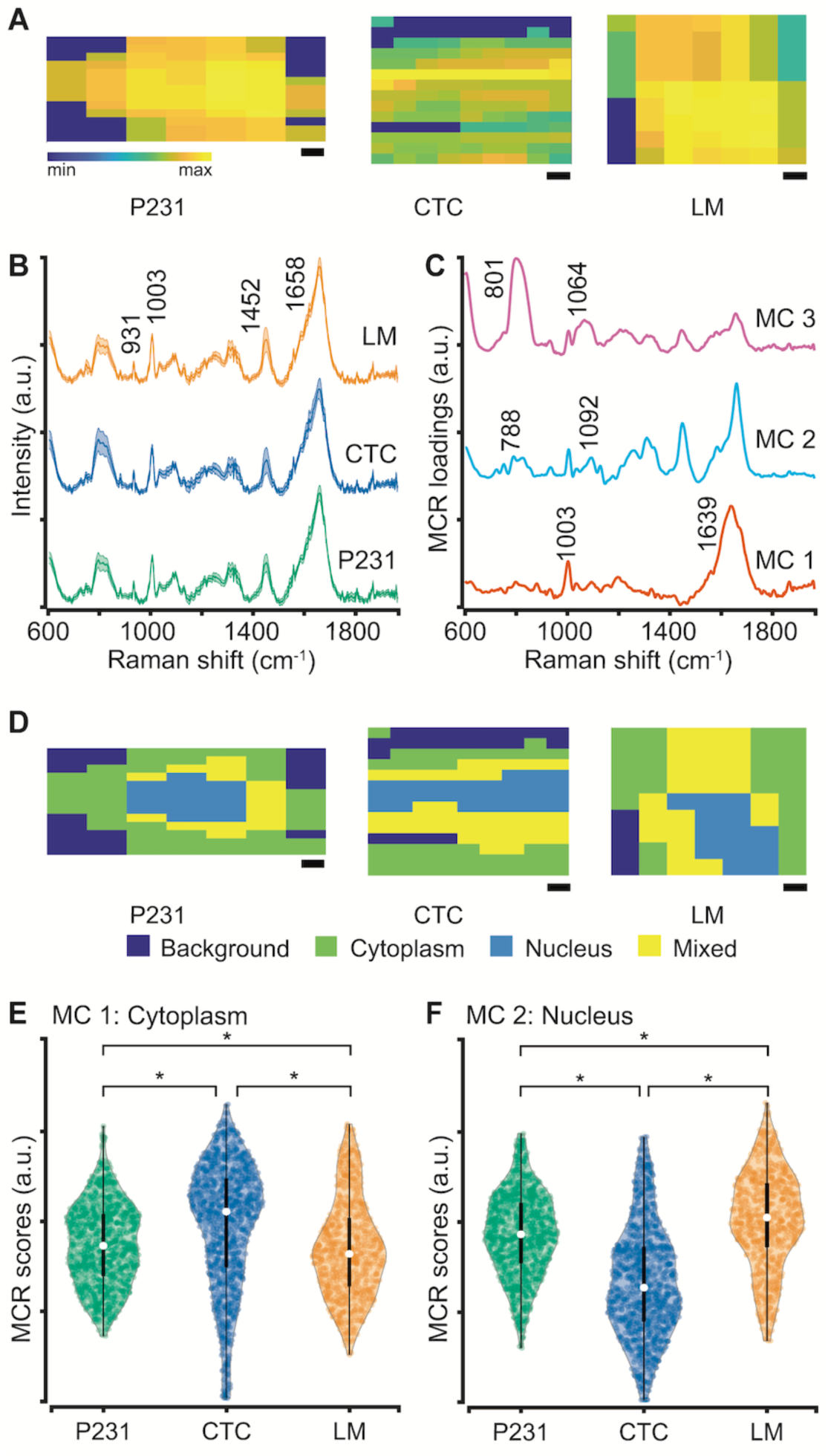
MCR segmentation of single-cell Raman images. **(A)** Representative coarse Raman maps reconstructed using the 1452 cm^-1^ peak intensity shown for P231, CTC, and LM cells. **(B)** Mean Raman spectra (with the shadow representing 1 s.d. and vertical offset for clarity) are shown and some prominent biological peaks highlighted for the three isogenic cell lines used in the study. **(C)** The three MCR component loadings derived from the combined spectral dataset are shown. MC1, MC2, and MC3 respectively show cytoplasm-like, nucleus-like, and quartz background spectral features. **(D)** The segmentation maps constructed by thresholding on MCR component scores for the cells in panel A are shown. The violin plots with embedded box and whisker plots show the distribution of MCR scores for cytoplasm-like **(E)** and nucleus-like **(F)** loadings. The scale bars represent 2 μm. * represents statistically significant differences at p < 0.05 threshold (Wilcoxon rank-sum test).

We further verified the assignment by reconstructing the abundance maps for the scores of MC1 and MC2. We assigned each pixel as cytoplasm, nucleus, or mixed by thresholding on the MC1 and MC2 scores. Using this MCR-ALS decomposition of spectral dataset allows better visualization of the spatial demarcation between cytoplasm and nucleus, which was not apparent in the coarse Raman maps at individual wavenumbers. The identification of pixels as those rich in cytoplasm and nucleus allow us to dissect the heterogeneous single-cell Raman measurements into relatively homogenous subsets for identifying subcellular compartments that capture the information necessary for identification of metastatic phenotypes. The remaining loading MC3, showing features at 801 cm^-1^ and 1064 cm^-1^, captures the minor contributions of quartz substrate in the cell spectra. We compared the scores of the cytoplasm-like and nucleus-like components to understand the relative abundance of these components in the cells of varying metastatic potential. The violin plots of MC1 and MC2 show that while the median values for the cytoplasm scores are significantly higher for the CTC in comparison to the P231 cells, the median for the LM cells is significantly lower in comparison to both P231 and CTC groups. Since the nucleus scores are negatively correlated with the cytoplasm scores, their medians show an opposite trend. The similarity of P231 and LM scores and their deviation from CTC hint at the ability of Raman spectroscopy to identify the differences associated with the EMT and MET processes that make CTC dissimilar to the P231 and LM cells. These observations are consistent with our prior characterization of these cell lines that showed significant differences in the mRNA expression levels of osteopontin (OPN), CD44, and vimentin (VIM) – genes involved in EMT, cell migration, extracellular matrix organization, and cell adhesion – in CTC and LM cells compared to the P231 cells [31]. Since the MCR scores represent the contribution of the pure component-like spectra to each spectrum irrespective of its location in the cell, the observed differences in the scores across the metastatic cascade provide relatively limited direct biological insights. However, these statistically significant differences in the MC component scores provide a rationale for exploring supervised classification techniques for the determination of metastatic phenotypes in the studied cancer cell lines.

We employed a multiclass random forest classifier to quantify our ability to classify P231, CTC, and LM cells based on the biochemical information encoded in their Raman spectra. Random forests are ensemble classifiers that employ a collection of decision trees constructed by random sampling of instances and variables in each tree to yield fast and generalizable models that are void of dependence on specific features or training instances [60]. Due to these characteristics and the ability to parallelize the tree construction, random forest classifiers are gaining attention in a variety of research areas including image classification and vibrational spectroscopy [61]. First, we subjected the entire spectral data consisting of spectra from all subcellular compartments (cytoplasm, nucleus, and mixed) to a leave-one-cell-out random forest classification task (as described in Methods). Briefly, we trained the classifier iteratively by leaving spectra from one cell at a time from the training dataset and subjecting it as a test dataset to the developed model. We observed satisfactory prediction performance across the three classes with only 4 misclassifications and 1 unclassification among 45 cells (**Fig. 4A**). The majority of misclassifications occurred between P231 and LM classes, while all CTC were classified accurately. This observation is in agreement with the similarity of both cytoplasm and nucleus MCR scores for P231 and LM classes and their deviation from CTC. The misclassifications of P231 cells can also be attributed to the presence of cells in the P231 group that have future propensity to intravasate into circulation and colonize lungs (i.e. future CTC and LM cells). Unlike most of the previous studies where the classification of spectra is done at a bulk level [23, 62, 63], the leave-one-cell-out analysis allowed us to demonstrate not only the ability to identify metastatic potential at a single-cell level but also the robustness of such classification by completely excluding representation of the test data from the training dataset.

**Figure 4.**
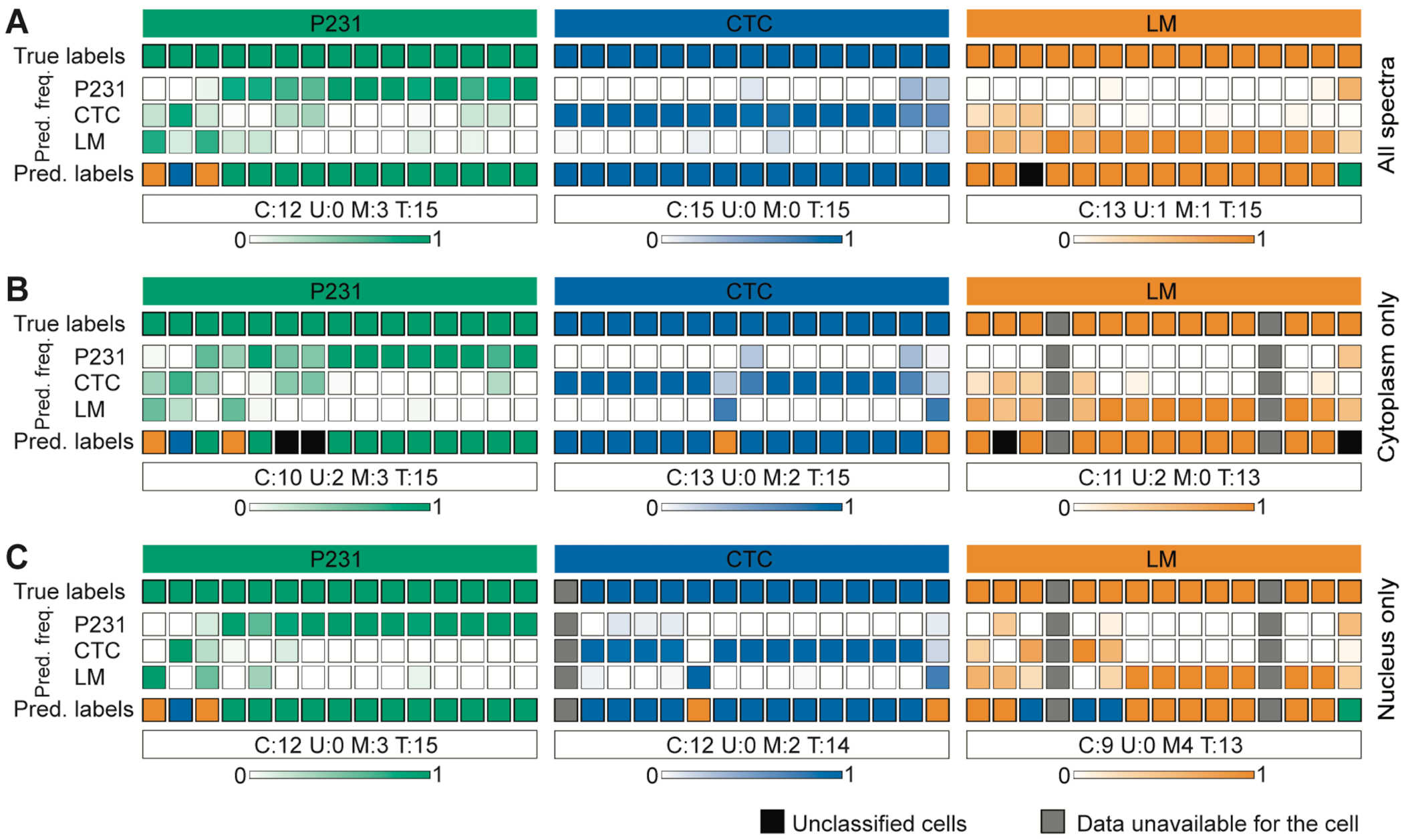
Leave-one-cell-out random forest classification of Raman images at a single-cell level. The leave-one-cell-out random forest predictions are shown for the multiclass classification task by including (**A**) all the spectra in the dataset, (**B**) spectra with high cytoplasm MC scores, and (**C**) spectra with high nucleus MC scores. Each column represents one unique cell, while the top and bottom rows respectively show the true and predicted class labels. The other rows show the normalized prediction frequencies (color bars at the bottom represent color scales) of spectra from each cell into the three classes. The classification results are summarized for each class to include the number of cells correctly classified (C), unclassified (U), and misclassified (M) out of the total (T) cells in the class.

While the leave-one-cell-out analysis provided an excellent prediction of phenotype for the cells in all the three classes, the use of the entire dataset comprised of spectra from different subcellular regions introduces substantial intra-class heterogeneity in the training dataset and may make prediction challenging. Such difficulty can be further exacerbated if the target phenotype changes are specifically guided by local molecular variations in particular regions, for example in the nucleus of cells treated with chemotherapeutic drugs. While there is no direct attribution of metastatic phenotypes observed in the isogenic panel employed in this study to specific compartments, we sought to train and test the random forest classifiers using subsets of the spectral dataset from the regions identified as cytoplasm and nucleus using MCR-ALS decomposition. The deviation of the observations from the baseline results obtained by subjecting the entire dataset will provide preliminary insights into specific localization of changes in subcellular regions that render the CTC and LM cells more metastatic in comparison to the parental P231 cells. First, we restricted our analysis to include only spectra that exhibit high scores for cytoplasm-like loading (MC1) in training and test datasets. The leave-one-cell-out analysis of the cytoplasm spectra (**Fig. 4B**) from the three classes yielded similar predictions with a slight improvement in the classification of LM cells, where a previously misclassified cell was now unclassified due to the relative increase of spectral classification into LM group. However, we found new unclassifications and misclassifications, respectively, in the P231 and CTC classes. Next, we performed the leave-one-cell-out analysis on the subset of dataset comprised only of spectra that show high scores of nucleus-like loading (MC2). We observed that the exclusion of cytoplasm spectra (**Fig. 4C**) resulted in the deterioration of performance in CTC and LM classes without affecting the P231 classification. Together, these results show that while the prediction of P231 and LM cells are primarily driven by the spectra acquired from the nucleus and cytoplasm respectively, the classification of CTC is more challenging and requires spectra from both regions. While these observations are preliminary and require further investigation in a larger cohort of cells, the results hint at the sufficiency of spectra from specific subcellular regions to predict subtle phenotypic differences associated with metastatic potential in closely related isogenic cells.

In conclusion, our label-free optical study revealed morphological and molecular differences among isogenic breast cancer cells of progressively increasing metastatic potential. Using 3D RI tomograms, we showed that the parental P231, circulating CTC, and lung metastatic LM cells showed subtle yet significant variations in morphology as assessed by area, aspect ratio, and cell dry mass. The observations were consistent with prior evidence of EMT and MET processes that guide the metastatic progression of these MDA-MB-231 breast cancer cells. To uniquely predict the metastatic potential of these cells with single-cell analytical resolution, we used Raman spectroscopic imaging to capture their biomolecular composition along with the spatial details. The use of MCR-ALS decomposition allowed better visualization and demarcation of the nucleus and cytoplasm despite the low resolution of the coarse Raman images. Finally, our random forest classification models incorporating a leave-one-cell-out strategy provided a route identification of subtle metastatic phenotype of cells at a single-cell level based on the coarse Raman maps. Further classification using the spectra individually from cytoplasm and nucleus regions as identified by MCR-ALS decomposition showed that specific subsets were sufficient for the identification of metastatic phenotypes. Taken together, these studies show that optical imaging and spectroscopy are sensitive to the differences in the cellular states guided by biological processes. We envision that coarse Raman imaging will be leveraged to build large spectral datasets from clinical samples that are amenable to machine learning analysis for determination of biomolecular phenotype/variant at a single-cell resolution to avoid loss of information associated with the population analyses. The imaging protocol and leave-one-cell-out random forest routine can readily be extended to the investigation of a variety of phenomena such as drug response, stem cell differentiation, and immune cell activation.

## Acknowledgments

S.K.P. acknowledges the support of the SLAS Graduate Education Fellowship Grant. I.B. acknowledges the support from the National Cancer Institute (R01 CA238025), the National Institute of Biomedical Imaging and Bioengineering (2-P41-EB015871-31) and the National Institute of General Medical Sciences (DP2GM128198). K.G. acknowledges the support from the National Cancer Institute (R01 CA213428, R01 CA213492). The authors thank Tomocube Inc for use of the 3D ODT system. The schematic in Figure 1A was partially created with BioRender.com.

